# NanoMethViz: an R/Bioconductor package for visualizing long-read methylation data

**DOI:** 10.1101/2021.01.18.426757

**Authors:** Shian Su, Quentin Gouil, Marnie E. Blewitt, Dianne Cook, Peter F. Hickey, Matthew E. Ritchie

## Abstract

**Motivation:** A key benefit of long-read nanopore sequencing technology is the ability to detect modified DNA bases, such as 5-methylcytosine. Tools for effective visualization of data generated by this platform to assess changes in methylation profiles between samples from different experimental groups remains a challenge.

**Results:** To make visualization of methylation changes more straightforward, we developed the R/Bioconductor package *NanoMethViz*. Our software can handle methylation calls generated from a range of different methylation callers and manages large datasets using a compressed data format. To fully explore the methylation patterns in a dataset, *NanoMethViz* allows plotting of data at various resolutions. At the sample-level, we use multidimensional scaling to look at the relationships between methylation profiles in an unsupervised way. We visualize methylation profiles of classes of features such as genes or CpG islands by scaling them to relative positions and aggregating their profiles. At the finest resolution, we visualize methylation patterns across individual reads along the genome using the *spaghetti plot,* allowing users to explore particular genes or genomic regions of interest.

In summary, our software makes the handling of methylation signal more convenient, expands upon the visualization options for nanopore data and works seamlessly with existing methylation analysis tools available in the Bioconductor project. Our software is available at https://bioconductor.org/packages/NanoMethViz.

## Introduction

Recent advances from Oxford Nanopore Technologies (ONT) have enabled high-throughput, genome-wide long-read DNA methylation profiling using nanopore sequencers, without the need for bisulfite conversion (1, 2).

A common goal of genome-wide profiling of DNA methylation is to discover differentially methylated regions (DMRs) between experimental groups. There is currently no software in the R/Bioconductor collection (3) for easily creating plots of methylation profiles in genomic regions of interest from the output of popular ONT-based methylation callers. We have developed *NanoMethViz* to create visualizations that give high resolution insights into the data to allow visual inspection of regions identified as differentially methylated by statistical methods. This software has been developed for compatibility with other software in the Bioconductor ecosystem (3), allowing for access to a wealth of existing statistical and genomic analysis methods. Specifically, this provides compatibility with the comprehensive toolkit for representing and manipulating genomic regions provided by *GenomicRanges* (4), and the statistical methods for DMR analysis available in packages such as *bsseq* (5), *DSS* (6) and *edgeR* (7).

The size of the data produced by ONT based methylation callers is the primary challenge in creating plots within defined genomic regions. It is not feasible to load entire methylation data-sets into memory on a standard computer, and for regions spanning the average length of a human or mouse gene, there are often enough data points to make smoothing visualizations computationally prohibitive. Together, this makes the analysis of methylation data difficult without access to high-performance computing (HPC), restricting the accessibility of methylation research using ONT sequencers.

## Design and Implementation

The *NanoMethViz* package provides conversion of data formats output by popular methylation callers *nanopolish* (5), *f5c* (8), and *Megalodon* into formats compatible with Bioconductor packages for DMR analysis.

At the time of writing, there is no consensus on the format for storing nanopore methylation data. The methylation callers *nanopolish, f5c* and *Megalodon* all produce slightly different outputs to represent similar information. Methylation calling from nanopore sequencing is still an active area of research and more formats are expected to arise. From the workflow presented in Figure 1A, *NanoMethViz* provides conversion functions from the output of various methylation callers into an intermediate format shown in Figure 1B, containing the minimal information for downstream processes. This intermediate format is used to create plots, and can be converted into various methylation count table formats and objects used by DMR detection functions using provided functions.

**Fig. 1.**
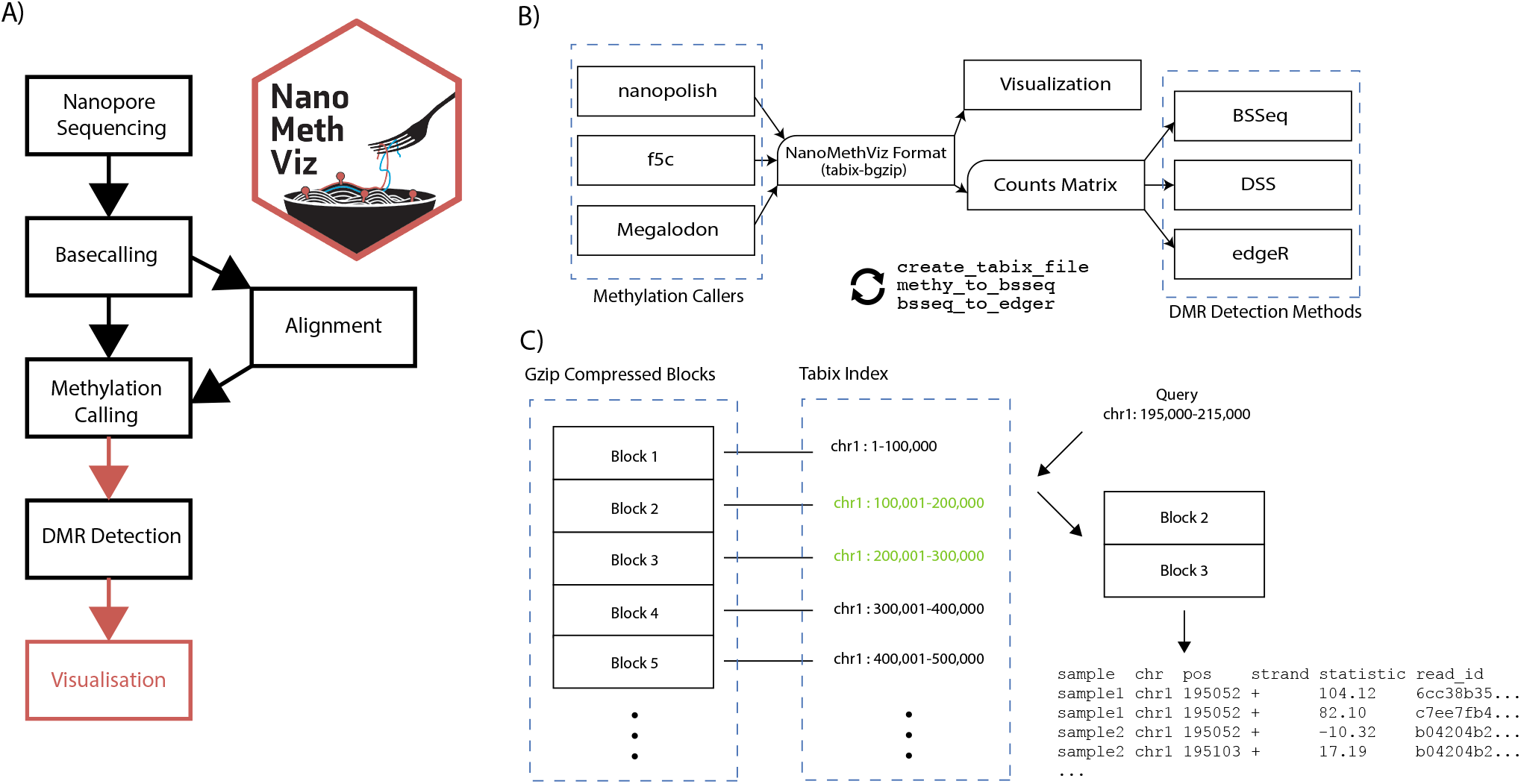
Nanopore methylation workflow and data format. A) The workflow used to perform differential methylation analysis. The red arrows indicate steps where further *NanoMethViz* provides conversion functions to bridge workflow steps. *NanoMethViz* performs visualization at the end of the workflow. B) Functions are provided in NanoMethViz to import the output of various methylation callers into a format used for visualization. This can be further converted by provided functions into formats suitable for various DMR detection methods provided in Bioconductor. C) The bgzip-tabix format compresses rows of tabular genomic information into blocks, and indexes the blocks with the range of genomic positions contained. This index is used for fast access the relevant blocks for decompression and reading.

*NanoMethViz* converts results from methylation caller into a tabular format containing the sample name, 1-based single nucleotide chromosome position, log-likelihood-ratio of methylation and read name. We choose log-likelihood of methylation as the statistic following the convention of *nanopolish*. This statistic can be converted to a methylation probability via the sigmoid transform as shown in Gigante *et al.* (2019) (9). The intermediate format and importing functions provided by *NanoMethViz* enables compatibility with existing methylation callers, as well as simplifying extension of support for future methylation caller formats. The information contained in this format is sufficient to perform genome wide methylation analysis as well as retain the molecule identities that are an advantage of long reads.

As shown in 1C, we compress the imported data using bgzip with tabix indexing. We use the tools *bgzip* and *tabix* included in *Rsamtools* toolkit (10, 11) to process the intermediate format; bgzip performs block-wise gzip compression such that individual blocks can be decompressed to retrieve data without decompressing the entire file, and tabix creates indices on position-sorted bgzip files to rapidly identify the blocks containing data within some genomic region. Having a format that is compressed with support for querying of data without loading in the whole data-set makes it feasible to analyse the data without the use of HPC, and allowing analysis to be performed on more widely available hardware.

Conversion is performed using block-wise streaming algorithms from the *readr* (12) package, this limits the amount of memory required to convert inputs of arbitrary size. Currently we support the import of methylation calls from *nanopolish*, *f5c* and *Megalodon*, and we also provide conversion functions from the tabix format into formats suitable for differentially methylated region analysis using *bsseq*, *DSS* or *edgeR* using `methy_to_bsseq` and `bsseq_to_edger`.

## Results

The primary plots provided by *NanoMethViz* are shown in Figure 2. They are the multidimensional scaling (MDS) plot for dimensionality reduced representation of differences in methylation profiles, the aggregate profile plot for methylation profiles of a set of features, and the *spaghetti plot* (9), for visualizing methylation profiles within specific genomic regions. While we have focused our development on 5mC methylation, in principle our work can be applied to any form of DNA or RNA modification.

**Fig. 2.**
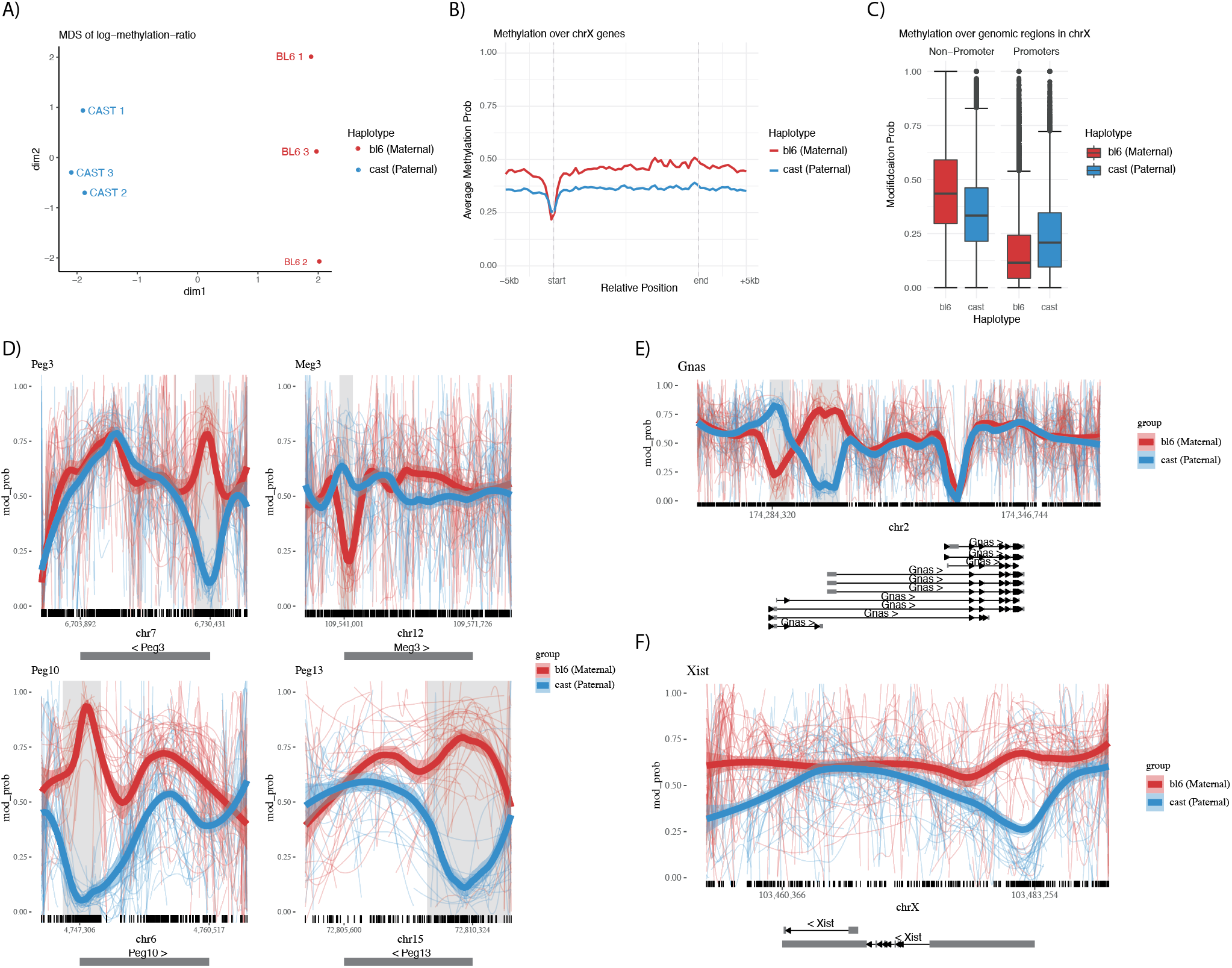
Summary of the plotting capabilities of *NanoMethViz.* A) Multidimensional scaling plot of haplotyped samples. B) Aggregated methylation profile across all genes in the X-chromosome, scaled to relative positions. C) Box plot of methylation probabilities over promoter and non-promoter regions for the BL6 and CAST haplotypes. D) Spaghetti plots of known imprinted genes Peg3, Meg3, Peg10 and Peg13. Thin lines show the smoothed methylation probability on individual long reads, the thick lines show aggregated trend across the all the reads. The shaded regions are annotated as DMR by *bsseq,* and the tick marks along the x-axis show the location of CpG motifs. E) Spaghetti plot of Gnas, which shows two adjacent regions of opposite imprinting patterns. F) Spaghetti plot of Xist. The transcription start site of the gene has visually apparently differential methylation but is not identified as such by *bsseq.*

We demonstrate the plots of *NanoMethViz* using a pilot dataset generated from triplicate female mouse placental tissues from F1 crosses between homozygous C57BL/6J mothers and CAST/EiJ fathers. Well characterized homozygous parents provided known SNPs for haplotyping reads (13), and the paternal X-chromosome is preferentially inactivated in female mouse placental tissue (14). Together these two properties allow the parent of origin and X-inactivation state of each read to be known a priori when performing analysis of methylation profiles. The three samples of E14.5 placental tissue were harvested and each sequenced using a single PromethION flow cell, the data was basecalled using *Guppy* (v3.2.2) using the `dna_r9.4.1_450bps_hac_prom.cfg` high accuracy profile. Reads were aligned to the GRCm38 primary assembly obtained from GENCODE (15), using *minimap2* (16) (v2.16) with the ONT profile set by `-x map-ont` argument. The output of *minimap2* was sorted and indexed using *samtools* (v1.9), and only primary alignments were retained for analysis. The retained reads were haplotyped using *WhatsHap* (17) using mouse variant information provided by the Sanger Institute (13). Methylation calling was performed by *f5c* (8) and associated with the haplotype information through the read IDs. *bsseq* (v1.22.0) (18) was used to identify differentially methylated regions and all visualisations in *NanoMethViz* were created using CRAN packages *ggplot2* (19) and *patchwork*.

The *MDS plot* shown in figure 2A is commonly used in differential expression analysis to summarize the differences between samples in terms of their expression profiles. It represents high dimensional data in lower dimensions while retaining the high dimensional similarity between samples. We use the log-methylation-ratio to represent the methylation profiles of samples and provide the conversion function `bsseq_to_log_methy_ratio` to convert from a `BSseq` object to a matrix of log-methylation ratios. This matrix can be used with the `plotMDS` function from the *limma* (20) Bioconductor package to compute MDS components for the most variable sites following the *edgeR* bisulphite sequencing analysis workflow (21). In Figure 2A, we see this approach shows separation of the haplotypes along the first dimension and according to sample (1,2,3) in the second dimension.

The *aggregation plot* shows aggregate methylation profiles across a class of features, revealing trends within a given class, such as promoters or repeat regions with fixed width flanking regions. It is produced by the function `plot_agg_regions`, which requires a table of genomic features or a `GRanges` object, and then plots the aggregate methylation profile scaled to the lengths of each feature such that they have the same start and end positions along the x-axis. The aggregation is an average of methylation profiles, with equal weights given to each feature as opposed to read, such that the aggregate is not biased towards features with higher coverage. This can be used to investigate specific classes of features such as genes or promoters. Figure 2B shows methylation profiles across all annotated genes in the X-chromosome, with the active X-chromosome (Xa) showing a higher level of methylation overall compared to the inactive X-chromosome (Xi). Genes from both chromosomes dip in methylation at the transcription start site, with Xi dipping below Xa by a small amount. This is further investigated in Figure 2C using the `query_methy` function to extract methylation data using ENSEMBL predicted promoters annotation to create a box plot. We see in the box plot higher levels of methylation in the maternal X-chromosome outside of promoter regions and lower levels of methylation within promoter regions. This matches previous observations in human fibroblast cells. (22)

The *spaghetti plot* created by the functions `plot_region` or `plot_gene` visualize the methylation profile of experimental groups within specific genomic regions. The plot shows methylation profiles of individual reads, annotations of CpG sites shown in tick marks along the x-axis, gene exons below the x-axis and top 500 most differentially methylated regions shaded in light grey. In 2D the well known family of Peg and Meg genes are shown, which are paternally expressed genes and maternally expressed genes, respectively. In the case of paternally expressed genes Peg3, Peg10 and Peg13, we see a drop in methylation in the paternal chromosomes near the TSS with an increase in methylation of the maternal chromosome. In the maternally expressed gene Meg3 we see a drop in methylation in the maternal chromosome but a relatively small increase in methylation in the paternal chromosome. Figure 2E shows the methylation profile of Gnas, with two oppositely imprinted regions adjacent to each other. Figure 2F shows the gene Xist, which is expressed from the inactive paternal X-chromosome, we can see reduced methylation near the TSS of the gene but it is not present in the top DMR results found by *bsseq.*

The *aggregate plots* and *spaghetti plots* both use `geom_smooth` from *ggplot2* to create smoothed methylation profiles. Of the smoothing methods provided by `geom_smooth`, we found *loess* gave the most aesthetically pleasing fits. However, we found that *loess* scales poorly with the number of data points typically found in this type of data. To resolve this, the *spaghetti plot* takes per-site means before calling `geom_smooth` to significantly improve performance. In the *aggregation plot*, the methylation profiles are aggregated across the features, with relative positions within feature bodies and the two fixed width flanking regions without scaling. It was found that the feature region tends to have a much higher density of data points than flanking regions, leading to poor smoothing behavior as *loess* selects *N* nearest points for fitting, with *N* being a fixed portion of the total data. Many more points from the model fitting will be taken from the feature region than the flanking regions near the boundary between feature and flanking regions. To overcome this issue, we take binned means along the relative genomic positions, which results in data of uniform density along the x-axis. These optimizations allow smoothed plots of the genomic regions or aggregate features to be created where it would otherwise be infeasible by naive usage of the `geom_smooth` function.

The features provided by *NanoMethViz* fill current gaps in the data flow between software in the nanopore methylation analysis pipeline. The performance focused implementation of the plotting allows them to be generated without the need of high performance computers, facilitating more accessible analysis.

Other major software for visualization of long-read methylation data includes Python packages *pycoMeth* and *Methplotlib*. *pycoMeth* provides a full workflow that produces a comprehensive interactive report on differentially methylated regions. *Methplotlib* is purely a plotting package for specified genomic regions.

Both *pycoMeth* and *Methplotlib* produce interactive plots of methylation data. *pycoMeth* produces summaries focused on CpG intervals, including a bar-plot with the count of methylation intervals, a heatmap of the methylation status of CpG intervals, density plot of the methylation log-likelihood of significant intervals, and a karyoplot of the density of significant CpG intervals along the chromosomes. It also provides a higher resolution heatmap and density plot for significant intervals. The significance testing uses the Mann-Whitney U test for two samples or Kruskal-Wallis H test for three or more samples, with Benjamini and Hochberg correction for multiple testing. *Methplotlib* creates detailed plots of specific genomic regions, including a line plot of the methylation frequencies of individual samples and a heatmap of the methylation profiles on individual reads.

Compared with *pycoMeth*, *NanoMethViz* does not provide a complete pipeline for analysis; rather it is intended to be used as a modular component of a workflow that includes other Bioconductor software for a more flexible and powerful analysis. *NanoMethViz* contains conversion functions to import data from methylation callers into its standard format, then conversions from the standard format into formats appropriate for DMR callers from Bioconductor, including *bsseq, DSS* and *edgeR.*

*Methplotlib* is similar in operation to *NanoMethViz* when plotting genomic regions. However, *Methplotlib* does not provide higher-level summaries such as the MDS plot or the feature aggregate plot. *NanoMethViz* also operates within interactive R sessions, as opposed to the command-line calls used by *Methplotlib,* which requires expensive reparsing of annotation files each time a new plot is created. *NanoMethViz,* therefore, better facilitates interactive exploration of long-read methylation data.

## Availability and Future Directions

The R/Bioconductor package *NanoMethViz* is available from https://bioconductor.org/packages/NanoMethViz. Vignettes are provided with examples of how to import data from methylation callers and how to create the basic plots. Example data is included with the package including data from genes Peg3, Meg3, Impact, Xist, Brca1 and Brca2. Data used for figures 2A, 2B and 2C can be found at https://zenodo.org/record/4495921.

In conclusion, *NanoMethViz* provides conversion functions, an efficient data storage format and a set of visualizations that allows the user to summarize their results at different resolutions. This work unlocks the potential for established Bioconductor DMR callers to be applied to data generated by ONT based methylation callers, lowers the hardware requirements for downstream analysis of the data, and provides key visualizations for understanding methylation patterns using ONT long reads.

Future development will support a wider range of plots, including some of those currently found in *pycoMeth* and *Methplotlib* to make them available for R users. Ongoing support will be added for any new, popular methylation callers that arise with differing formats to existing callers.

## Acknowledgments

We thank Kathleen Zeglinski for designing the *NanoMethViz* logo and Kelsey Breslin and Tamara Beck for their assistance in generating the data used to test our software. This work was supported by Australian National Health and Medical Research Council (NHMRC) Project grant 1098290 to MER and MEB, a Bellberry-Viertel Senior Medical Research Fellowship to MEB, Victorian State Government Operational Infrastructure Support and Australian Government NHMRC IRIISS.

## Bibliography

1. Jacob Schreiber, Zachary L Wescoe, Robin Abu-Shumays, John T Vivian, Baldandorj Baatar, Kevin Karplus, and Mark Akeson. Error rates for nanopore discrimination among cytosine, methylcytosine, and hydroxymethylcytosine along individual DNA strands. Proc. Natl. Acad. Sci. U. S.A.,110(47):18910–18915, November 2013.

2. Andrew H Laszlo, Ian M Derrington, Henry Brinkerhoff, Kyle W Langford, Ian C Nova, Jenny Mae Samson, Joshua J Bartlett, Mikhail Pavlenok, and Jens H Gundlach. Detection and mapping of 5-methylcytosine and 5-hydroxymethylcytosine with nanopore MspA. Proc. Natl.Acad. Sci. U. S.A.,110(47):18904–18909, November 2013.

3. Robert C Gentleman, Vincent J Carey, Douglas M Bates, Ben Bolstad, Marcel Dettling, Sandrine Dudoit, Byron Ellis, Laurent Gautier, Yongchao Ge, and Jeff Gentry Bioconductor: open software development for computational biology and bioinformatics. Genome Biol., 5 (10):R80, 2004.

4. Michael Lawrence, Wolfgang Huber, Hervé Pagès, Patrick Aboyoun, Marc Carlson, Robert Gentleman, Martin T Morgan, and Vincent J Carey. Software for computing and annotating genomic ranges. PLoS Comput. Biol., 9(8):e1003118, August 2013.

5. Jared T Simpson, Rachael E Workman, P C Zuzarte, Matei David, L J Dursi, and Winston Timp. Detecting DNA cytosine methylation using nanopore sequencing. Nat. Methods, 14 (4):407–410, 2017.

6. Yongseok Park and Hao Wu. Differential methylation analysis for BS-seq data under general experimental design. Bioinformatics, 32(10):1446–1453, 2016.

7. Mark D Robinson, Davis J McCarthy, and Gordon K Smyth. edgeR: a Bioconductor package for differential expression analysis of digital gene expression data. Bioinformatics, 26(1): 139–140, 2010.

8. Hasindu Gamaarachchi, Chun Wai Lam, Gihan Jayatilaka, Hiruna Samarakoon, and Martin A Smith. GPU Accelerated Adaptive Banded Event Alignment for Rapid Comparative Nanopore Signal Analysis. bioRxiv Bioinformatics, pages 1–37, 2019.

9. Scott Gigante, Quentin Gouil, Alexis Lucattini, Andrew Keniry, Tamara Beck, Matthew Tinning, Lavinia Gordon, Chris Woodruff, Terence P Speed, Marnie E Blewitt, and Matthew E Ritchie. Using long-read sequencing to detect imprinted DNA methylation. Nucleic Acids Res., 47(8):e46, 2019.

10. Martin Morgan, Hervé Pagès, Valerie Obenchain, and Nathaniel Hayden. Rsamtools: Binary alignment (BAM), FASTA, variant call (BCF), and tabix file import, 2020. R package version 2.4.0.

11. Heng Li. Tabix: Fast retrieval of sequence features from generic TAB-delimited files. Bioinformatics, 27(5):718–719, March 2011.

12. Hadley Wickham, Jim Hester, and Romain Francois. readr: Read Rectangular Text Data, 2018. R package version 1.3.1.

13. Thomas M Keane, Leo Goodstadt, Petr Danecek, Michael A White, Kim Wong, Binnaz Yalcin, Andreas Heger, Avigail Agam, Guy Slater, Martin Goodson, Nicholas A Furlotte, Eleazar Eskin, Christoffer Nellåker, Helen Whitley, James Cleak, Deborah Janowitz, Polinka Hernandez-Pliego, Andrew Edwards, T Grant Belgard, Peter L Oliver, Rebecca E McIntyre, Amarjit Bhomra, Jérôme Nicod, Xiangchao Gan, Wei Yuan, Louise Van Der Weyden, Charles A Steward, Sendu Bala, Jim Stalker, Richard Mott, Richard Durbin, Ian J Jackson, Anne Czechanski, José Guerra-Assunçao, Leah Rae Donahue, Laura G Reinholdt, Bret A Payseur, Chris P Ponting, Ewan Birney, Jonathan Flint, and David J Adams. Mouse genomic variation and its effect on phenotypes and gene regulation. Nature, 477(7364): 289–294, September 2011.

14. N Takagi and M Sasaki. Preferential inactivation of the paternally derived X chromosome in the extraembryonic membranes of the mouse. Nature, 256(5519):640–642, August 1975.

15. Jennifer Harrow, Adam Frankish, Jose M Gonzalez, Electra Tapanari, Mark Diekhans, Felix Kokocinski, Bronwen L Aken, Daniel Barrell, Amonida Zadissa, Stephen Searle, If Barnes, Alexandra Bignell, Veronika Boychenko, Toby Hunt, Mike Kay, Gaurab Mukherjee, Jeena Rajan, Gloria Despacio-Reyes, Gary Saunders, Charles Steward, Rachel Harte, Michael Lin, Cédric Howald, Andrea Tanzer, Thomas Derrien, Jacqueline Chrast, Nathalie Walters, Suganthi Balasubramanian, Baikang Pei, Michael Tress, Jose Manuel Rodriguez, Iakes Ezkurdia, Jeltje Van Baren, Michael Brent, David Haussler, Manolis Kellis, Alfonso Valencia, Alexandre Reymond, Mark Gerstein, Roderic Guigó, and Tim J Hubbard. GENCODE: The reference human genome annotation for the ENCODE project. Genome Res., 22(9):1760–1774, 2012.

16. Heng Li. Minimap2: Pairwise alignment for nucleotide sequences. Bioinformatics, 34(18): 3094–3100, 2018.

17. Marcel Martin, Murray Patterson, Shilpa Garg, Sarah O Fischer, Nadia Pisanti, Gunnar W Klau, Alexander Schöenhuth, and Tobias Marschall. WhatsHap: fast and accurate read-based phasing. bioRxiv, page 085050, 2016.

18. Kasper D Hansen, Benjamin Langmead, and Rafael A Irizarry. BSmooth: from whole genome bisulfite sequencing reads to differentially methylated regions. Genome Biol., 13 (10), 2012.

19. Hadley Wickham. ggplot2: Elegant Graphics for Data Analysis. Springer-Verlag New York, 2016. ISBN 978-3-319-24277-4.

20. Matthew E Ritchie, Belinda Phipson, Di Wu, Yifang Hu, Charity W Law, Wei Shi, and Gordon K Smyth. Limma powers differential expression analyses for RNA-sequencing and microarray studies. Nucleic Acids Res., 43(7):e47, 2015.

21. Yunshun Chen, Bhupinder Pal, Jane E Visvader, and Gordon K Smyth. Differential methylation analysis of reduced representation bisulfite sequencing experiments using edgeR. F1000Res., 6:2055, 2018.

22. Michael Weber, Jonathan J Davies, David Wittig, Edward J Oakeley, Michael Haase, Wan L Lam, and Dirk Schübeler. Chromosome-wide and promoter-specific analyses identify sites of differential DNA methylation in normal and transformed human cells. Nat. Genet., 37(8): 853–862, August 2005.

